# Structural and functional characterization of SidF, a possible dual substrate *Aspergillus fumigatus* N5-acetyl-N5-hydroxy-L-ornithine transacetylase

**DOI:** 10.1101/2024.08.03.606473

**Authors:** Thanalai Poonsiri, Nicola Demitri, Jan Stransky, Hubertus Haas, Michele Cianci, Stefano Benini

**Author notes:** Dept. of Pathobiology, Faculty of Science, Mahidol University, Rama VI Road, Ratchathewi, Bangkok 10400 THAILAND.

## Abstract

Siderophore-mediated iron acquisition is essential for the virulence of *Aspergillus fumigatus*, a fungus causing life-threatening aspergillosis. Developing drugs targeting the siderophore biosynthetic pathway could help improve disease management. The transacetylases SidF and SidL generate intermediates for different siderophores in *A. fumigatus*. *A. fumigatus* has a yet unidentified transacetylase that complements SidL during iron deficiency in SidL-lacking mutants.

We present the first X-ray structure of SidF, revealing a conserved two-domain architecture with tetrameric assembly. Importantly, the N-terminal domain contributes to protein solubility and oligomerization, while the C-terminal domain containing the GCN5-related N-acetyltransferase (GNAT) motif is crucial for the enzymatic activity and mediates oligomer formation. Notably, AlphaFold modelling demonstrated structural similarity between SidF and SidL. Enzymatic assays showed that SidF can utilize acetyl-CoA as a donor, previously thought to be a substrate of SidL but not SidF, and selectively uses N5-hydroxy-L-ornithine as an acceptor. Based on these findings, we propose SidF as the unknown transacetylase complementing SidL activity, highlighting its central role in *A. fumigatus* siderophore biosynthesis.

This study elucidates the structure of SidF and reveals a novel role in siderophore biosynthesis. Investigation of this uncharacterized GNAT protein enhances our understanding of fungal virulence and holds promise for its potential application in developing antifungal therapies.

## Introduction

*Aspergillus fumigatus* is a ubiquitous saprophytic fungus living on organic debris. Nevertheless, it is also an opportunistic pathogen that can cause life-threatening invasive pulmonary aspergillosis [1]. While the incidence is higher among immunocompromised patients and those with underlying pulmonary diseases, the population at risk is steadily expanding, also due to SARS-CoV2 infections [2–4]. Additionally, evidence is emerging that resistance to triazole antifungals, the predominant class of drugs used to treat fungal diseases [5], is increasing worldwide [4]. This trend underscores the importance of monitoring *Aspergillus* as a potential drug-resistant threat.

Iron, with its redox properties, is a vital cofactor for essential biochemical reactions in diverse life forms [6]. However, strict homeostasis is necessary to balance its potential toxicity [6–8]. The low environmental bioavailability of iron in the environment, driven by its propensity to form insoluble ferric (Fe^3+^) hydroxides, necessitates specific acquisition mechanisms, particularly for pathogens exploiting host iron stores [6, 9]. *A. fumigatus* employs two main iron uptake strategies: low-affinity ferrous (Fe^2+^) iron and high-affinity ferric iron uptake systems including siderophore-mediated iron acquisition (SIA) and reductive iron assimilation (RIA) [10, 11]. Siderophores are low molecular mass, ferric iron-specific chelators [10] and this research focuses on SIA, a key virulence factor during host infection [12–14]. *A. fumigatus* produces two fusarinine-type siderophores, termed fusarinine C (FsC) and triacetylfusarinine C (TAFC), and two ferrichrome-type siderophores, termed ferricrocin (FC) and hydroxyferricrocin (HFC) [10, 15]. FsC TAFC, and FC are secreted to capture environmental iron [10, 12, 15, 16]; FC is also employed for intracellular iron handling within hyphae, and HFC is employed for conidial iron storage [10, 16, 17].

Both fusarinine- and ferrichrome-type siderophores share the same starting point, the conversion of ornithine to N5-hydroxyornithine by the monooxygenase SidA [14], but then the pathways diverge. A scheme of the *A. fumigatus* siderophore biosynthesis pathway is shown in Fig. 1. Ferrichrome biosynthesis utilizes the transacetylase SidL and an additional uncharacterized enzyme for N5-acetylation [18], followed by assembling FC from serine, glycine, and N5-acetyl-N5-hydroxyornithine, catalyzed by the nonribosomal peptide synthetase (NRPS) SidC [16]. An unknown enzyme hydroxylates FC to hydroxyferricrocin. Fusarinine-type siderophore synthesis involves mevalonate conversion to anhydromevalonyl-CoA by the mevalonyl-CoA ligase SidI and the mevalonyl-CoA hydratase SidH [19]. Anhydromevalonyl-CoA is transferred to N5-hydroxyornithine by the transacylase SidF to yield N5-anhydromevalonyl-N5-hydroxyornithine [16]. The NRPS SidD then links three N5-anhydromevalonyl-N5-hydroxyornithine units to form FsC, which SidG subsequently acetylates to yield TAFC [6, 10, 20, 21]. GCN5-related N-acetyltransferases (GNATs), an ubiquitous superfamily, encompasses enzymes across all life domains, crucial for diverse processes from bacterial antibiotic resistance to circadian rhythms [22]. Typically, the enzyme acylates the primary amine of an acceptor molecule using a donor molecule such as acetyl-CoA (AcCoA). Their conserved fold consists of β-sheets and α-helices (β1-H1-H2-β2-β3-β4-H3-β5-H4-β6) with a characteristic conserved V-shaped cleft for acyl-CoA binding (β4-β5) and a variable acceptor substrate binding site reflecting GNAT functional diversity [23]. The catalytic site is placed between the donor and acceptor sites [24].

**Figure 1.**
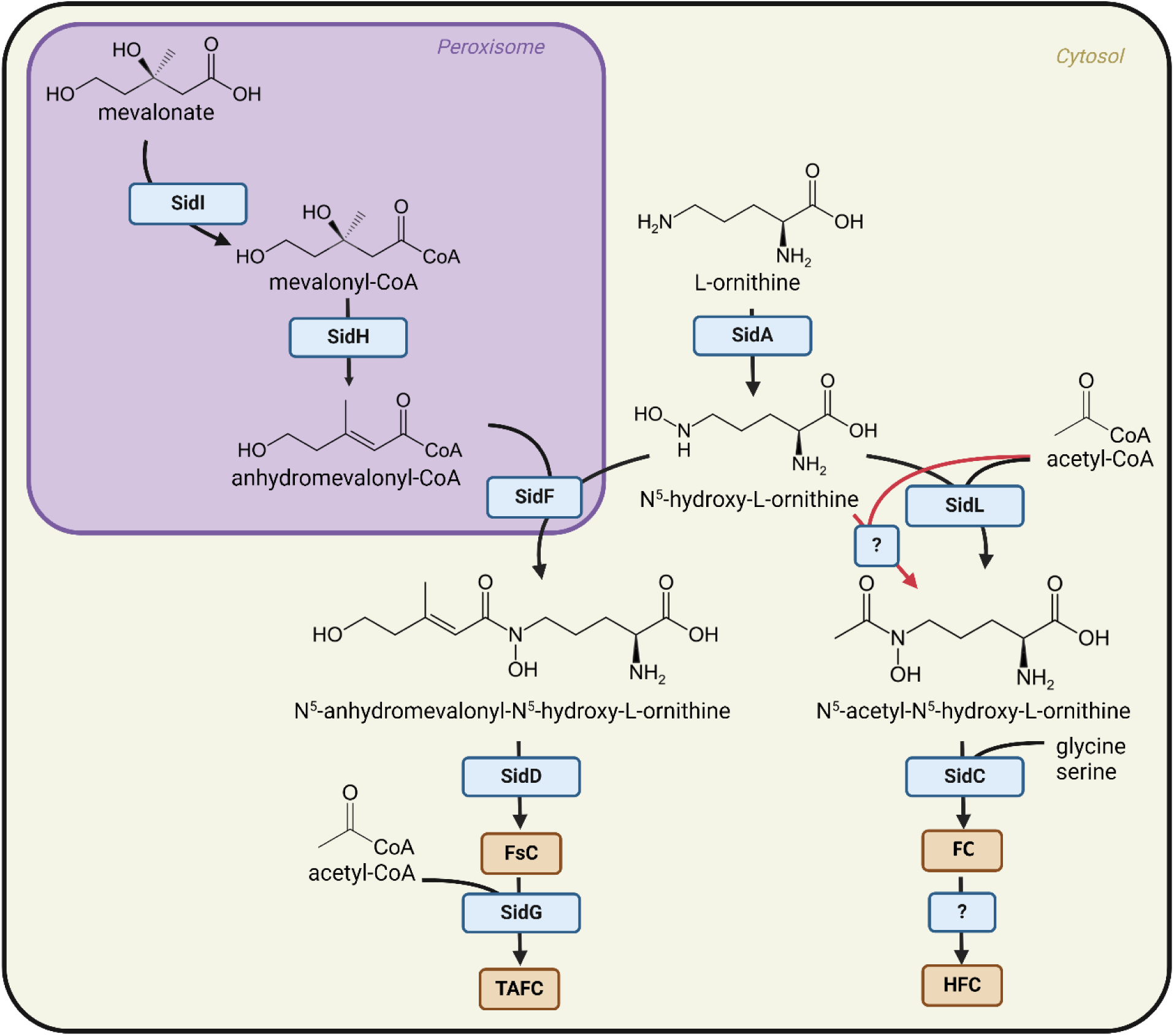
*A. fumigatus* siderophore biosynthetic pathway. Involved enzymes are boxed in blue and described in the text. Siderphores are boxed in brown. Red arrows denote reactions involving unknown enzyme. Peroxisome and cytoplasm are shaded in purple and yellow, respectively. Adapted from “Plant cell with organelles”, by BioRender.com (2024). Retrieved from https://app.biorender.com/biorender-templates. Created with BioRender.com. / Mahidol University

According to the InterPro database, *A. fumigatus* (ATCC MYA-4609) has 47 GNAT type proteins that belong to different protein families (pfam). Notably, SidF and SidL are the only two *Aspergillus* GNATs classified within the pfam13523 (EC number 2.3.1.-), which exhibits a broad taxonomic distribution, predominantly bacterial (86.9%) with a smaller fungal representation (12.2%). Despite sharing the same functional class, SidF and SidL display significant differences [10, 18]. The SidF encoding gene is on chromosome 3 clustered with other siderophore biosynthesis genes, while the SidL encoding gene is on chromosome 1. SidF, is localized in the peroxisome and coregulated with other siderophore genes by SreA in response to iron levels, distinct from cytosolic SidL, whose regulation appears to be largely independent of iron availability or SreA [18, 25]. Moreover, SidF and SidL share only limited amino acid sequence similarity in their C-terminal domains [18, 25]. Interestingly, *A. fumigatus* possesses an unidentified N5-acetyl-N5-hydroxy-L-ornithine transacetylase activity that complements SidL function during iron limitation, although its molecular identity remains unknown [18].

Though many GNATs are intensively studied, most are still unexplored. Given the established role of siderophore biosynthesis as a crucial virulence factor in *A. fumigatus*, targeting enzymes within this pathway holds promise for novel antifungal drug development. SidF has unique characteristics and warrants further investigation, despite sharing functional similarities with SidL. This study integrates protein structural analysis with enzymatic assays to comprehensively characterize SidF.

Our study revealing the high-resolution structure and biochemical characterization of SidF enabled us to propose that SidF might complement SidL during iron starvation, opening new avenues for therapeutic intervention against *A. fumigatus* infections.

## Materials and Methods

### Plasmid Construction

A synthetic SidF DNA construct (UniProt Q4WF55), optimized for Escherichia coli expression, was obtained from Genscript and cloned into the NdeI/BamHI restriction sites of the pET-28a(+)-TEV vector. The recombinant protein was expressed with an N-terminal His-tag. The theoretical molecular weight and isoelectric point of SidF were calculated to be 55.7 kDa and 6.54, respectively. To generate N- and C-terminal deletion mutants, the “around the horn” PCR method [26] was employed using primers listed in Supplementary Table S1.

### Protein Expression and Purification

SidF expression was induced by either autoinduction or 1 mM IPTG at 30°C overnight in *E. coli* BL21(DE3). Lysis buffer was 50 mM Tris pH 8.0, 500 mM NaCl, 0.5 mM ethylenediaminetetraacetic acid (EDTA), 10 mM β-mercaptoethanol (BME), and 20 mM imidazole. Cell lysis was performed by sonication. The sample was centrifuged at 12000xg for 30 minutes at 4°C and then purified by using Ni-NTA in 50 mM Tris pH 8, 200 mM NaCl, and 20 mM imidazole binding buffer. Then, washed and eluted with the same buffer containing 20 mM imidazole and 250 mM imidazole, respectively. Elution fraction was loaded onto a Superdex™ 200 Increase 10/300 GL Tricorn™ size-exclusion chromatography (SEC) column (GE HealthCare) equilibrated with 50 mM Tris pH 8.0, 200 mM NaCl at a flow rate of 0.5 mL·min^-1^. The peak at a retention volume of 12.18 mL was concentrated and used for crystallization.

SidF N-terminal domain and SidF C-trim were expressed and purified with the same procedure. Except that the SEC column for the N-terminal domain was Superdex™ 75 Increase 10/300 GL Tricorn™.

### Crystallization, Diffraction Experiment, Data Processing and Model Building

SidF crystallization conditions were searched using commercial screens from Molecular Dimension and Hampton Research at a protein concentration of 10 mg/ml by the micro batch-under the volatile oil method at 20 °C. Crystals obtained from 0.2 M potassium thiocyanate, and 20% polyethylene glycol (PEG) 3350 were used for diffraction experiments. The crystals were flash-frozen in liquid N_2_. Cryoprotection was achieved by supplementing the reservoir solution with 12% glycerol. X-ray data were collected at a cryogenic temperature (100K) at the ELETTRA 11.2C beamline, Elettra Synchrotron Trieste, Italy [27]. Data were reduced by XDS [28] and scaled by Aimless [29]. The predicted three-dimensional structure, generated by The ColabFold AlphaFold2 [30, 31], was used as a template for molecular replacement *via* MOLREP [32]. The process succeeded with R/Rfree=0.25/0.30. The structure was refined by REFMAC5 [33] and built in COOT [34] in CCP4Cloud [35] or CCP4i2 [36]. The final model was refined with a weight term of 0.5 with resolution limits of 1.87 Å. Data collection and refinement statistics are shown in Table 1. Models were visualized using Chimera [37] and ChimeraX [38].

**Table 1.** Diffraction data collection and refinement statistics.

| Data collection | SidF (7QF6) | SidF Nter (8KD8) |
| --- | --- | --- |
| Space group | *P* 1 | *C* 222<sub>1</sub> |
| Cell dimensions |  |  |
| a, b, c (Å) | 80.27 80.84 179.84 | 62.2 110.67 69.47 |
| α, β, γ(°) | 101.168 91.668 117.252 | 90 90 90 |
| Resolution (Å) | 174.82-1.87(1.90-1.87) | 29.42 – 2.58(2.70-2.58) |
| R<sub>merge</sub> | 0.061(0.459) | 0.095(0.6) |
| R<sub>pim</sub> | 0.038(0.290) | 0.04(0.255) |
| I/σI | 17(3.6) | 18.1(4.4) |
| CC half | 0.999(0.929) | 0.998(0.942) |
| Completeness (%) | 97.1 (96.2) | 99.8(99.2) |
| Redundancy | 6.8(6.8) | 12.1(12.3) |
| **Refinement** |  |  |
| Resolution (Å) | 174.82-1.87 | 29.42 – 2.58 |
| Total No. reflections | 4925967 | 95327 |
| R<sub>work</sub> / R<sub>free</sub> | 0.17/0.21 | 0.17/ 0.24 |
| No. atoms | 30901 | 2712 |
| Protein | 28812 | 2681 |
| Ions | 2 | - |
| ligands | 114 | - |
| Waters | 1973 | 31 |
| B-factors |  |  |
| Protein | 33.37 | 53.88 |
| Ions | 60.21 | - |
| ligands | 43.07 | - |
| Waters | 34.79 | 48.41 |
| R.M.S. deviations |  |  |
| Bond lengths (Å) | 0.015 | 0.013 |
| Bond angles (°) | 1.972 | 2.192 |

SidF N-terminal domain and SidF C-trim were purified as the full-length protein. SidF N-terminal crystals were obtained from 0.2 M potassium sodium tartrate tetrahydrate, 20 % w/v polyethylene glycol 3,350 and subsequently cryoprotected with 40% Morpheus® precipitant mix 3. The structure was solved by molecular replacement using MrBUMP [39]. SidF C-trim crystals were obtained from 0.1 M Sodium cacodylate pH 6.5, 40% v/v MPD, 5% w/v PEG 8000 but no X-ray diffraction was observed. For SidF C-trim, X-ray diffraction experiment was performed at a cryogenic temperature at beamline P13 operated by EMBL Hamburg at the PETRA III storage ring (DESY, Hamburg, Germany) [40].

### Computational tools

A conserved domain search was performed using the National Center for Biotechnology Information (NCBI) tool [41] (https://www.ncbi.nlm.nih.gov/Structure/cdd/wrpsb.cgi). The Basic Local Alignment Search Tool (BLAST) was used to compare protein sequences to sequence databases [42]. Protein structures database search was retrieved from the Protein Data Bank (PDB) (https://www.rcsb.org). Protein structure comparison was conducted using the PDBeFold service at European Bioinformatics Institute [43] (http://www.ebi.ac.uk/msd-srv/ssm), RCSB PDB tool (pairwise structure alignment) (https://www.rcsb.org/alignment), or US-align (multiple structure alignment) [44] (https://zhanggroup.org/US-align/). Analysis of macromolecular interfaces was done using ‘Protein Interfaces, Surfaces and Assemblies’ service PISA at the European Bioinformatics Institute [45] (http://www.ebi.ac.uk/pdbe/prot_int/pistart.html). The computed models of SidF and SidL were generated by the web-based service AlphaFold2 (40) (https://colab.research.google.com/github/sokrypton/ColabFold/blob/main/AlphaFold2.ipynb).

### Small Angle X-ray Scattering (SAXS)

SAXS was measured using a laboratory SAXS instrument (SAXSpoint 2.0, Anton Paar) at IBT CAS (Vestec, Czech Republic); the instrument was equipped with X-ray source MetalJet C2 (Excillum) and detector Eiger R 1M. The X-ray wavelength was 1.34 Å.

SidF N-terminal domain and SidF C-terminal domain were measured using a batch mode: sample (5mg/ml in 50 mM Tris-HCl, 200 mM NaCl pH 8) was loaded to thin wall quartz capillary (diameter of 1.5 mm) and exposed for 15 min with sample to detector distance of 571 mm. The corresponding buffer was measured under the same condition. Scattering empty capillaries and capillaries filled with water were measured for 5 min to perform absolute scale calibration.

SidF full-length was analyzed with SEC-SAXS mode: the SAXS system was connected to a fast protein liquid chromatography (FPLC) system AKTAGo (GE Healthcare. Samples were loaded onto a Superdex™ 200 Increase 10/300 GL (GE HealthCare) at 20°C; the separation was performed with flow a rate of 0.7 ml/min which was lowered to 0.05 ml/min upon peak detection with absorbance at 280 nm in AktaGO. Data were collected at a distance of 791 mm. The measurement cell allows in-situ measurement of UV-Vis absorption spectra (CaryUV 60, Agilent technologies) which were continuously measured in a range from 200 nm to 400 nm with a repetition rate of 30s.

Data processing was conducted in PRIMUS [41] and custom software. SEC-SAXS chromatographs were analyzed using RAW [46]. The molecular mass was calculated from Bayesian Inference. Comparison of scattering profiles was done in Crysol [42] and FoXS [43]. The *ab initio* model included averages from 15 independent model calculations with symmetry (P222) or without symmetry (P1) determined using DAMMIF [44]. The model data were averaged with DAMAVER [45] and refined with DAMMIN [46]. The low-resolution-model surface representation was created in CHIMERA using the ‘molmap’ command. Atomic models were validated using CRYSOL and FoXS. Final model of C-trim was calculated using CORAL [47]. EOM [48] was further used for validation of the flexible loops: 10000 models were of the flexible N-terminus was generated using RANCH [48], and the most probable ensamble selected using GAJOE [48]. Essential SAXS data acquisition, sample details, data analysis, and modelling fitting are given in Supplementary table S2.

### Broad-Substrate Screen for Gcn5-Related N-Acetyltransferases

The broad-substrate screen was adapted from M. Kuhn et al, 2013 [49]. The experiments were conducted in conventional 96-well PCR plates or tubes in a reaction volume of 50 µL. The reaction consisted of 50 mM Tris pH 8.0, 0.5 mM AcCoA, and 5 mM substrates. A wide variety of acceptors were used including cadaverine, L-lysine, L-glutamic acid, glycine, kanamycin A, ampicillin, chloramphenicol, nicotinamide adenine dinucleotide (NAD), urea, guanidine HCl, L-ornithine, oxidized glutathione (GSSH), hydroxylamine, and N5-hydroxy-L-ornithine. Cadaverine was freshly prepared before use. The reaction was initiated by adding 0.06 µM freshly prepared enzyme diluted in 50 mM Tris pH 8.0 and 200 mM NaCl. The reaction was carried out for 8 minutes at 25°C before being stopped with 50 µL of 100 mM Tris pH 8.0 and 6 M guanidine HCl. For free thiols detection, 200 µL of developer (0.2 mM 5,5’-dithiobis (2-nitrobenzoic acid) (DTNB), 100 mM Tris pH 8.0, and 1 mM EDTA) was immediately added and incubated for 10 min at room temperature. 290 µL of reaction was transferred to 96-well clear polystyrene flat-bottom plate and the absorbance was measured at 405 nm with 595 nm background subtraction in a Tecan Infinite F200 microplate reader. Each reaction had its control which was the identical condition without enzyme. All assays were performed in duplicate for each protein purification batch. A standard curve was created by preparing 2-fold dilutions of N-acetyl-L-cysteine ranging from 500 to 3.9 µM and measuring in duplicate for each independent assay.

## Results and Discussion

### 1. SidF structural characterization

#### The SidF monomer comprises two domains, with the N-terminal domain resembling the C-terminal domain

The SidF monomer contains 18 β-strands and 14 helices (both 3-10 and α-helix) divided into an N-terminal (residues 1–191) and a C-terminal domain (residues 227–462) (Fig. 2). There is no electron density for residues 1–24. The SidF N-terminal domain is composed of 8 β-strands (1–8) that form a barrel-like structure, wrapping around helix 3 (H3) and fringed with loops and helixes between β-strands (Fig. 2). Similarly, the C-terminal domain shows a barrel-like structure containing 10 β-strands wrapping around helix 11 (H11). The gaps of the half-barrel structure in each domain are facing each other. The two domains are connected by a long loop.

**Figure 2.**
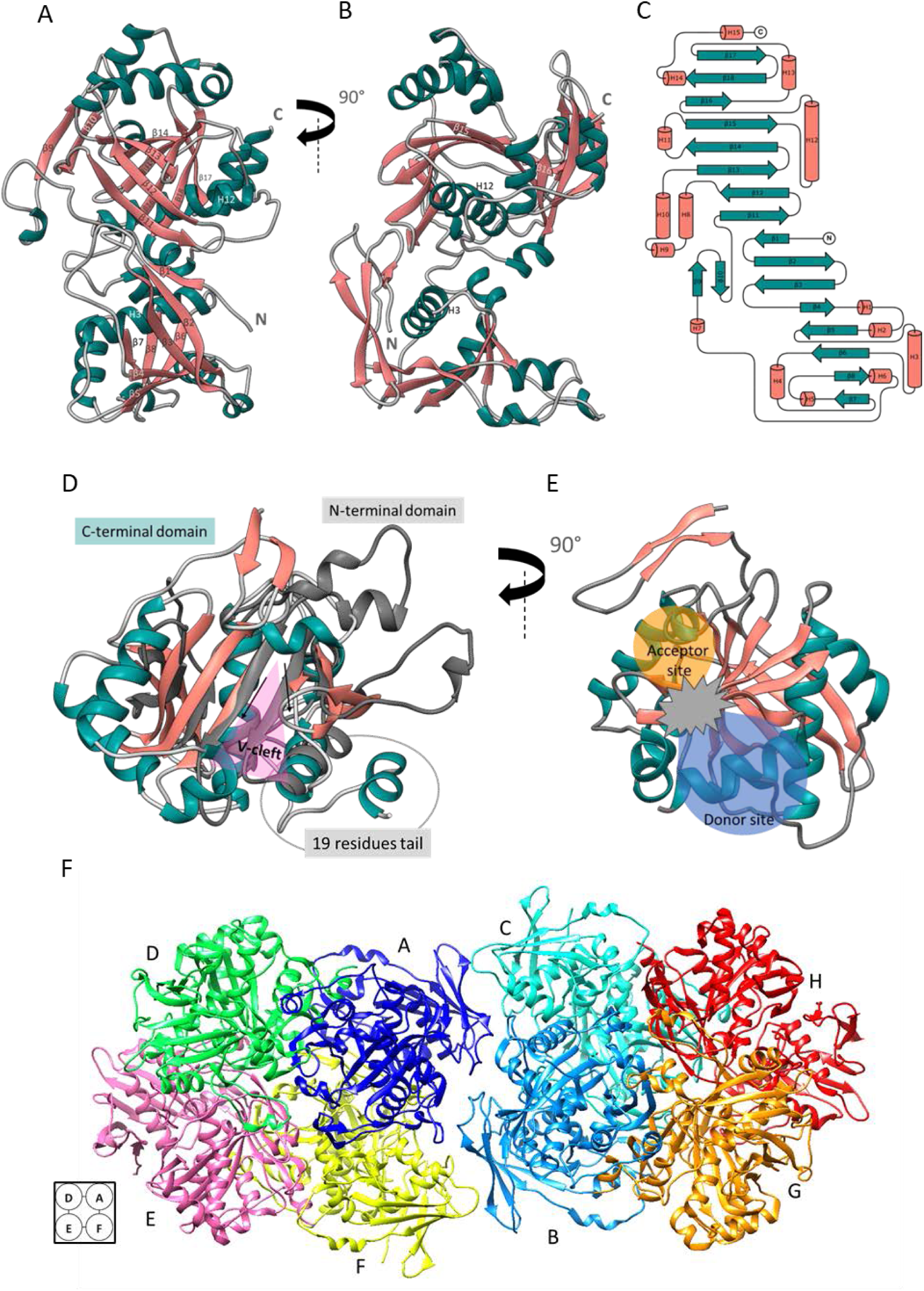
Crystal structures of SidF. (A) Ribbon representation of a SidF monomer (PDB entry 7QF6). Helices are colored green and labeled with “H”, β-strands are colored salmon and labeled with “β”, and loop regions are colored grey. (B) The same SidF monomer rotated 90° counter-clockwise along the vertical axis. (C) Topology diagram of SidF monomer generated with TopDraw [56]. β represent the β-strand and H represent helix (D) Superposition of the N-terminal and C-terminal domains of SidF. The C-terminal domain (residues 200-462) is colored identically to Figure 1A, while the N-terminal domain is shown in dark grey. The superposition highlights the absence of the V-cleft donor binding site in the N-terminal domain. The 19-residue C-terminal tail is circled. (E) The C-terminal domain of SidF rotated 90° counter-clockwise along the vertical axis. This view reveals the characteristic GNAT fold with the putative acceptor, donor, and catalytic sites clearly visible. (F) Ribbon representation of the asymmetric unit of SidF crystallized in the P1 space group, illustrating eight protein chains (colored uniquely by chain) assembled into two tetramers. The inset on the left depicts the arrangement of individual subunits within a tetramer.

Searching the PDB with the search term “Gcn5-related N-acetyltransferases,” gives 105 protein structures representative of each UniProt accession with an Enzyme Commission number (EC number) 2.3.1.- (accessed on 23 October 2023). Of these, 73.3% are small (calculated mass 12.8–31.8 kDa) and contain only the GNAT motif. SidF has a mass of about 53.3 kDa, larger than typical GNAT proteins, and consists of two structural similar N- and C-terminal domains. It was previously reported that the GNAT N-myristoyl transferase monomer harbors two GNAT domains with an internal two-fold symmetry. Despite the poor sequence identity of the two domains, this arrangement was proposed to be the outcome of gene duplication [50]. This could possibly be the case for SidF.

However, the amino acid sequence search using NCBI conserved domain tools identified the conserved GNAT motif only in the SidF C-terminal domain and not in the N-terminal domain. The protein BLAST (BLASTP) identified many homologs of SidF with coverage of 93-100% encoded by several other *Aspergillus spp.* and various ascomycetes. A Position-Specific Iterated BLAST (PSI-BLAST) search identified a wider range of distantly related homologs from ascomycetes and γ-proteobacteria, albeit with reduced coverage (40–87%). These lower-coverage homologs aligned primarily to the C-terminal region of SidF, encompassing the conserved GNAT domain. Moreover, the N-terminus of SidF is not conserved outside the phylum Ascomycota.

By using PDBeFold (accessed on 4 January 2023), the structure similarity search agreed with the sequence search, and the conserved acetyltransferase at the C-terminal domain of SidF matched with several acetyltransferase proteins. In another search, the N-terminal domain alone gave the same results, suggesting a similarity between the two domains. Superimposition of SidF C-terminal domain onto DfoC [51] from *Erwinia amylovora* (PDB ID: 5O7O) yielded a low sequence identity (30%) but a high structural conservation (RMSD 1.274 Å). Similar results were obtained when comparing SidF C-terminal domain to human N-alpha-acetyltransferase 80 [52] (PDB ID: 6NBE, 12% identity, 2.66 Å RMSD).

To further explore the similarity between the N- and C-terminal domains, the SidF model was bisected at residue 200 and the two domains were superimposed (Fig. 2D). Pairwise structure alignment showed an RMSD of 4.28 and a Template Modeling (TM)-score of 0.44, indicating that the two protein structures compared are different both locally and globally. Though the comparison gave a TM-score below the cut-off of 0.5, we noted that the N-terminal domain is not completely dissimilar to the C-terminal domain. The main difference was the absence of the substrate-binding V-cleft in the SidF N-terminal domain (Fig. 2D), which explains the absence of the GNAT motif in the N-terminal domain and the poor structure alignment score.

#### SidF is observed as a unique tetramer in the crystal structure and in solution

Numerous enzymes within the GNAT superfamily are known to exist as oligomers. While the majority of GNATs are typically observed in dimeric states, it’s worth mentioning that examples of monomeric, trimeric, tetrameric, hexameric, and even dodecameric configurations have also been documented [23]. Moreover, the GNAT domain can also be part of a large protein with multiple domains [51]. The structure of SidF differs from that of other GNAT proteins, which form dimers from each small monomer with a continuous β-sheet [22], an inserted β-sheet [53], or a helical bundle at the dimer interface [54]. The SidF monomer has two similar (TM score 0.44) independent folding units in the N- and C-terminal halves which are connected by a long loop (Fig 2). Eight chains (two tetramers) of SidF are observed in the asymmetric unit of the P1 symmetry crystal (Fig. 2F). According to PISA, SidF chains A and D engage extensively, forming a dimer with 43 hydrogen bonds and 31 salt bridges, yielding a high Complex Formation Significance Score (CSS) of 0.946 [45]. Similarly, SidF chains E and F create another dimer, featuring 43 hydrogen bonds and 29 salt bridges, with a CSS of 0.946. The interaction extends further as the adjacent dimers link chains A to F and D to E to form a tetramer. SidF chains A and F are stabilized by 31 hydrogen bonds, 15 salt bridges, with a CSS of 0.727. Additionally, SidF chains D to E form an interaction characterized by 33 hydrogen bonds, 16 salt bridges, with a CSS of 0.727. These inter-chain interfaces collectively contribute to the robustness and stability of the tetramer assembly (no interaction between chain AE and DF) (Fig. 2F). However, the tetramer-tetramer interaction is not significant with a CSS value of 0.

The SidF molecular mass of 157 kDa was estimated by SEC corresponds to a lighter mass than the tetramer, as compared to standard proteins.

SAXS studies confirmed the tetrameric nature of SidF in solution (Fig. 3 and Supplementary Fig. S2). The radius of gyration (Rg) determined through Guinier analysis was 40.3 ± 0.3 Å. The pair distribution function (P(r)) indicated features consistent with a prolate particle, revealing a maximum intra-particle distance (Dmax) of 190 Å. The molecular mass estimation for SidF, derived from Bayesian Inference, yielded a value of 208 kDa corresponding to SidF tetrameric oligomerization. Nevertheless, the maximum dimension does not align with the measurement observed in the X-ray structure, which is only 97.9 Å (Supplementary Fig. S3). The 1-24 N-terminal residues are disordered as suggested by low Alphafold model confidence score and confirmed by the X-ray structure. Thus, there might be variations in these flexible ends that affect the Dmax measurement.

**Figure 3.**
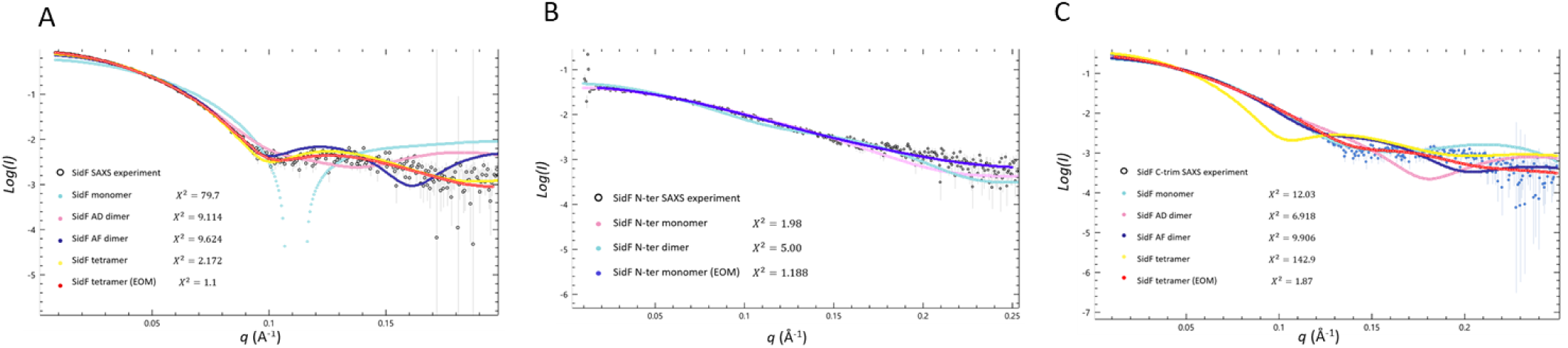
The comparative analysis of the calculated SAXS profiles of the SidF crystal structure and the experimental SAXS data for SidF (A) SidF wild-type. Experimental SAXS scattering curve is shown in blank scattering with error bars, while the scattering profile of SidF monomer, dimers, tetramer, and AlphaFold model were calculated with CRYSOL are shown in different colors. The scattering profile of SidF tetramer calculated with EOM shown in dark blue. Chi-square value is indicated. (B) SidF N-terminal domain. Experimental SAXS scattering curve is shown in blank scattering with error bars, while the scattering profile of SidF N-terminal domain monomer, and dimers calculated with FoXS are shown in light blue and pink, respectively. The scattering profile of SidF N-terminal domain monomer calculated with EOM shown in dark blue. Chi-square value is indicated. (C) SidF C-trim. Experimental SAXS scattering curve is shown in blank scattering with error bars, while the scattering profile of SidF monomer, dimers, and tetramer model were calculated with CRYSOL are shown in different colors. The scattering profile of SidF tetramer calculated with CORAL shown in dark blue. Chi-square value is indicated.

The comparative analysis involved aligning the SAXS profiles derived from the SidF crystal structure with the experimental SAXS data for SidF (Fig. 3). The tetrameric SidF structure exhibited a satisfactory fit to the experimental data, yielding a chi-square value of 2.172 and ruling out the possibility of monomeric or dimeric arrangements. EOM was used to take the flexible loop into account, which resulted in perfect fit (chi-square of 1.1), where the compact core is identical with the crystal structure.

#### Structure analysis of SidF N-terminal domain

The full length of the SidF X-ray structure shows discrete N- and C-terminal domains. The two domains are connected by a loop and form 12 hydrogen bonds, with no part of the N-terminal domain intervening in the C-terminal domain. To further explore the function of SidF N- and C-terminal domains, the SidF full-length was divided into two domains at residue 200 (Supplementary Fig. S4), and each one was expressed and purified under the conditions used for the full-length protein. SidF N-terminus (residue 1-199) was successfully expressed and purified, while the expressed SidF C-terminal domain (residue 200-462) was insoluble. Both protein constructs were expressed with an N-terminal His-tag.

The crystal structure of the SidF N-terminal domain was solved as a monomer in *C222_1_* space group at a maximum resolution of 2.58 Å. Its monomer displayed the same topology observed in the full-length protein, except that the residues 25-49 extended outward instead of being tucked in (Fig. 4). The N-terminal His-tag and the residues 1-24 were not visible in the electron density maps. Upon generating symmetry mates, the extended N-terminal region interacted with the corresponding region from another monomer, resulting in a stable dimer with a CSS score of 1 (according to PISA).

**Figure 4.**
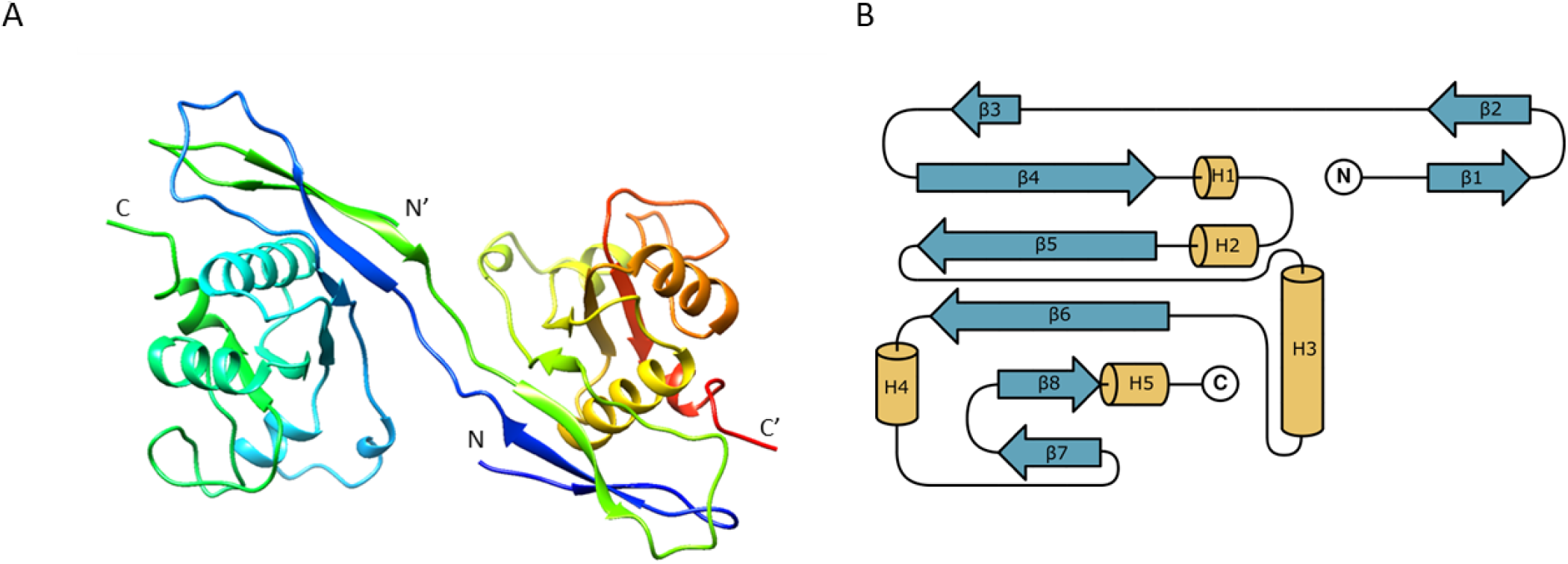
(A) Ribbon representation of the SidF N-terminal domain dimer generated from crystal symmetry (PDB entry 8KD8). (B) Topology diagram of SidF N-terminal domain monomer generated with TopDraw [56]. β represent the β-strand and H represent helix.

While several nearby symmetry mates were produced, no interactions were predicted (CSS score = 0), indicating a lack of stability and suggesting the possibility of crystal contact artifacts rather than biological contacts. Besides, the protein was eluted from size exclusion chromatography column with approximate mass of 53.84 kDa supporting the formation of a stable dimer.

To confirm the dimeric nature of the SidF N-terminal domain, SAXS investigations were conducted (Supplementary Fig. S5). Unexpectedly, SAXS analysis did not support dimer conformation. The Guinier analysis determined a radius of gyration (Rg) of 21.85 ± 0.33 Å. The pair distribution function (P(r)) revealed characteristics resembling a prolate particle, with a maximum intra-particle distance (Dmax) of 90 Å. Bayesian Inference estimated the molecular mass as 21.2 kDa, corresponding to the monomer rather than the dimer seen in the crystal structure.

Comparing the calculated SAXS profiles from the dimeric crystal structure of the SidF N-terminal domain to the SAXS experimental data (Fig. 3) resulted in a poor fit with a chi-square value of 5.00. However, using the monomer structure significantly improved the fit, yielding a chi-square value of 1.98. Further refinement using the EOM led to a further improvement in the fit, with a chi-square value of 1.19. PISA analysis of the SidF structure indicates that the N-terminal domain possesses a propensity to exist as a monomer in solution. The formation of the X-ray quaternary structure is primarily driven by hydrogen bonding interactions of the C-terminal domain, with the N-terminal domain potentially contributing to inter-chain interactions with the C-terminal domain of other chains. These findings suggest that the N-terminal domain alone lacks the essential structural elements required for stable dimer or tetramer formation in solution.

In the crystal structure, the dimer of the SidF N-terminal domain shows a similar fold to that of full-length SidF. This phenomenon, characterized by the partial exchange of N-terminal residues between monomers, resulting in the formation of a continuous β-sheet, bears resemblance to the established dimerization interface observed in typical GNAT family proteins [23]. However, the biological significance of this dimeric form remains uncertain, as it lacks evidence from the SAXS analysis. The observed N-terminal crystallographic dimerization of SidF might be attributed to crystal packing. Further studies are needed to experimentally validate these findings.

#### Structure analysis of SidF C-trim

Attempts to isolate the C-terminal domain in a soluble form were unsuccessful. To improve C-terminal solubility, we employed a domain truncation strategy targeting predicted disordered regions at both the N- and C-termini of the C-terminal domain, using structural data from full length SidF and AlphaFold modelling (Supplementary Fig. S4). Despite attempts to generate soluble C-terminal truncates for further characterization, none were successful. Notably, a construct encompassing the full-length protein with a C-terminal truncation of 19 amino acids (residues 1-443), designated SidF C-trim, remained soluble.

The SidF C-trim protein exhibited comparable solubility to the full-length version. SEC analysis suggested a molecular mass of approximately 206.25 kDa similar to the wild-type protein and potentially with the same tetrameric assembly. However, subsequent SAXS data challenged this hypothesis (Supplementary Fig. S6).

Guinier analysis of the SAXS data revealed a radius of gyration (Rg) of 34.52 ± 0.2 Å. The pair-distribution function (P(r)) indicated a prolate ellipsoidal particle with a maximum dimension (Dmax) of 150 Å. Bayesian Inference estimated the molecular mass of the C-trim construct to be 109.1 kDa, consistent with dimeric oligomerization.

Comparative analysis of the experimental SAXS data with the calculated dimeric forms derived from the SidF X-ray crystal structure AD and AF dimers yielded chi-square values of 6.918 and 9.906, respectively (Fig. 3). These relatively high chi-square values suggest a potential discrepancy between the solution-state conformation of the C-trim dimer and the dimeric interfaces observed in the crystal structure. Employing CORAL, which combines rigid body modelling with ab-intio modelling, resulted in a new arrangement of the SidF dimer.

Analysis of the full-length structure revealed the presence of 14 hydrogen bonds within the 19 C-terminal residues of the AD dimer interface, whereas no such hydrogen bonds were observed in the equivalent region of the AF dimer. This suggests that the C-terminal 19 residues may be crucial for the formation of the AD dimer.

As a result, it is plausible that, beyond its unidentified function, the SidF N-terminal domain plays a crucial role in ensuring the solubility of the C-terminal domain. The C-terminal domain is crucial for tetrameric formation. Without the C-terminal domain, the N-terminal domain alone appears as a monomer. Moreover, the 19-residue tail contributes to tetramer formation through interactions between dimers.

### 2. SidF has most likely N5-acetyl-N5-hydroxy-L-ornithine transacetylase activity

#### *In Vitro* Analysis of SidF’s substrate specificity

Although SidF and SidL are different in many aspects such as gene location, gene regulation, and protein subcellular location, a BLASTP sequences search demonstrated significant similarity between SidF and SidL, with an alignment score of 154, 61% coverage, 31.63% percent identity, and an E-value of 6.6 x 10^-46^. Moreover, they are classified in the same Pfam, have similar functions and protein folding. SidF catalyzes the transfer of an anhydromevalonyl group from anhydromevalonyl-CoA to the N5-hydroxyornithine primary amine. Similarly, SidL transfer an acetyl group from acetyl-CoA to the N5-hydroxyornithine primary amine. The AlphaFold model of SidL is similar to SidF with RMSD of 3.14 Å, reference coverage of 74%, and TM-score of 0.76 with sequence identity of only 29% (Fig. 5). This supports the hypothesis that SidF might compensate for SidL in the knockout experiment [18].

**Figure 5.**
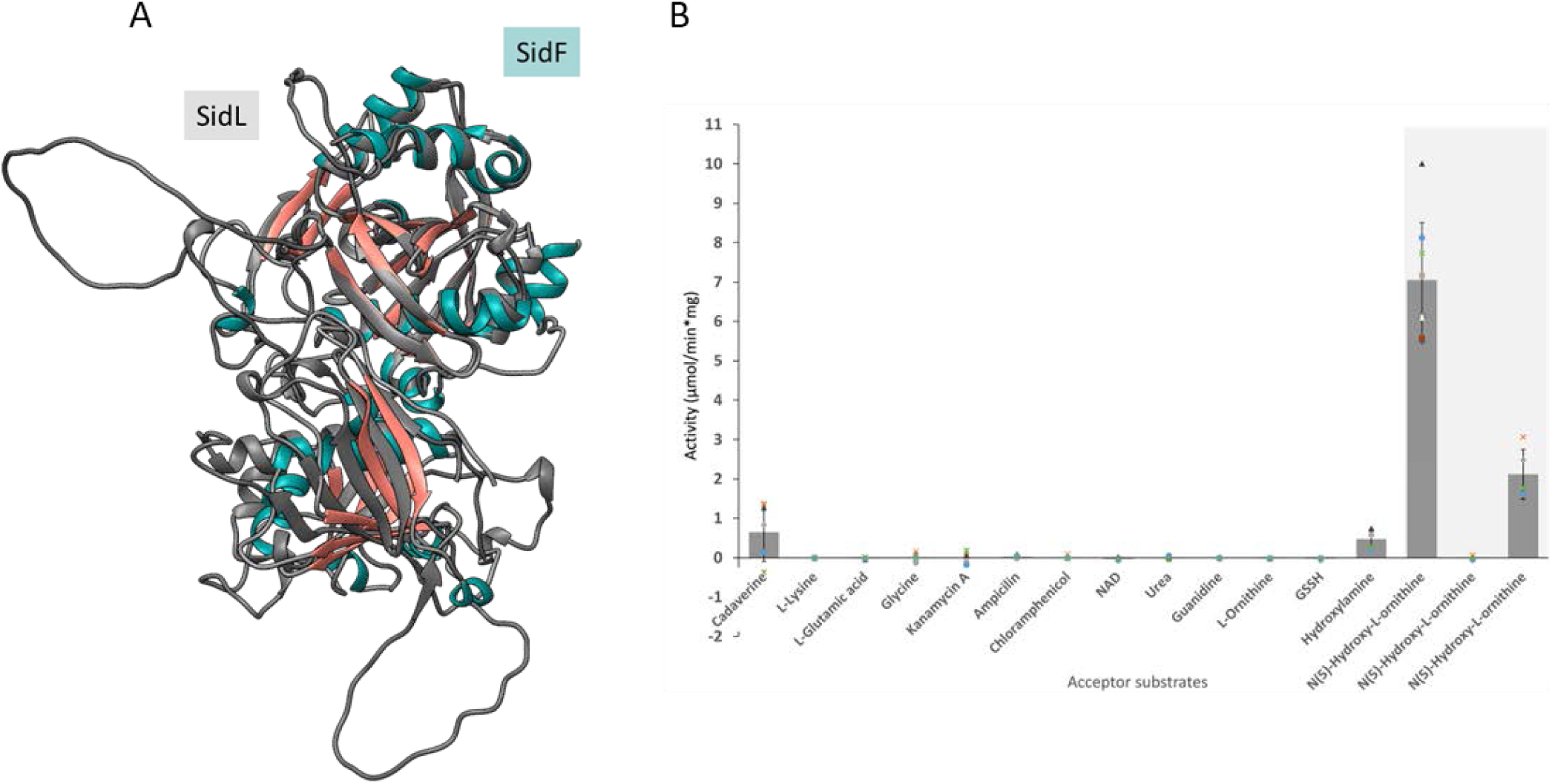
(A) Superposition of the SidF crystal structure and SidL AlphaFold model (B) DTNB assay The SidF full-length protein was used to evaluate substrate utility (column 1-14). The assay was setup with AcCOA donor and variety of acceptor substrates, where NAD is nicotinamide adenine dinucleotide and GSSH is oxidized glutathione. Highlighted in grey are the reactions setup to test activity of the SidF variants, SidF full-length, SidF N-terminal domain (N) and SidF C-trim, with AcCOA donor and N5-hydroxy-L-ornithine acceptor substrates. The average activity is presented along with the standard deviation, as well as individual data points (n=5, except for column 14, where n=10).

To test this hypothesis, in vitro assays were conducted to assess the acetyltransferase activity of SidF using acetyl-CoA, the preferred donor substrate for SidL, and N5-hydroxyornithine acceptor. The DTNB assay was employed to detect the exposed thiol groups following acetyl transfer reaction. The results confirmed the capacity of SidF to utilize acetyl-CoA as a donor (Fig. 5). This finding supports the hypothesis that SidF might compensate for SidL in knockout experiments, further investigation would elucidate the compensation mechanism *in vivo*.

### 3. Functional Characterization of SidF

#### GNAT Broad-Substrate Screen

GNAT family enzymes exhibit a generally conserved donor site and a more variable acceptor site. This versatility allows them to acetylate diverse metabolites, antibiotics, and proteins, potentially leading to multiple functionalities [49]. To evaluate the substrate range of SidF, a DTNB assay was employed with AcCoA as the donor substrate and a variety of potential acceptor substrates (Fig. 5). SidF exhibited no activity on L-ornithine, but activity on the natural substrate N5-hydroxy-L-ornithine notably increased to 7.05 ± 1.45 µmol/min/mg (p-value < 0.0005). The sole distinction between L-ornithine and N5-hydroxy-L-ornithine is the presence of the hydroxyl group, which suggests that the hydroxyl group may play a significant role in substrate recognition. To further investigate, hydroxylamine, which contains a hydroxyl group (OH) and an ammonia group (NH_2_), was used to assess the impact of the hydroxyl group. SidF demonstrated the ability to process hydroxylamine, but with a relatively low activity of 0.48 ± 0.23 µmol/min/mg (p-value = 0.001936). This suggests that the hydroxyl group influences SidF acceptor recognition to a limited extent. SidF may employ a substrate recognition mode analogous to MbtK/IucB-like proteins (InterPro entry IPR019432). 82.6% of the proteins of this group are bacterial and specifically recognize hydroxyl groups [55]. Cadaverine showed a detectable signal as acceptor substrate (0.65 ± 0.74 µmol/min/mg), but the high standard deviation rendered the difference from background non-significant. Conversely, SidF did not utilize lysine, glutamic acid, glycine, kanamycin A, ampicillin, chloramphenicol, NAD, urea, guanidine, or GSSH as acceptor substrates, highlighting its substrate specificity.

#### Functional Characterization of SidF N-Terminal Domain and C-Trim

DTNB assay employing AcCoA as the donor substrate and N5-hydroxyornithine acceptor was used to assess the enzymatic activity of the SidF N-terminal domain and SidF C-trim. The N-terminal domain exhibited no detectable activity compared to the background control (p-value = 0.575804) (Fig. 5). This observation aligns with the predicted absence of enzymatic activity within the N-terminus due to the lack of the conserved GNAT motif.

The dimeric form of SidF C-trim, displayed a significant reduction (69.95%) in activity compared to the full-length protein (p-value = 0.000012). This observation underscores the importance of tetramerization for optimal catalytic efficiency of SidF. Furthermore, the C-terminal tail’s proximity to the acyl-CoA binding site suggests a potential regulatory role by influencing the accessibility of the donor substrate.

#### Conclusions

This study presents a comprehensive investigation of the structure and function of *A. fumigatus* SidF. X-ray crystallography, SAXS analysis, and enzyme activity assay support the biological relevance of the tetrameric assembly of the full-length protein. The SidF subunits possess a repetitive structure, resulting in a size larger than that typically observed for GNAT family enzymes. This characteristic is common to homologs from other *Aspergillus* species and numerous other ascomycetes. Notably, SidF structure represents the first structural characterization of this group. The absence of GNAT activity in the SidF N-terminal domain, consistent with the lack of a conserved GNAT motif, raises questions about its functional necessity. Our findings suggest that this region might be essential for solubility and proper tetramer assembly. The C-terminal domain is essential for tetramer formation with its 19-residue tail which may specifically contribute to the dimerization. Our analysis further suggests that SidF recognizes the hydroxyl group in the acceptor substrate. Based on structural similarities and shared donor substrate utilization, SidF is proposed to be a functional counterpart of SidL.

Our results provide insights into the structure of SidF and propose a novel function for SidF in siderophore biosynthesis. The study of this uncharacterized GNAT protein enhances our understanding of *A. fumigatus* siderophore biosynthesis and has promising implications for disease treatment.

## CRediT authorship contribution statement

**Thanalai Poonsiri**: Conceptualization, Methodology, Investigation, Writing – original draft. **Nicola Demitri**: Methodology, Investigation. **Jan Stransky**: Methodology, Investigation. **Michele Cianci**: Methodology, Investigation. **Hubertus Haas**: Conceptualization, Supervision. **Stefano Benini**: Conceptualization, Methodology, Investigation, Funding acquisition, Supervision. All contributing authors participated in the manuscript development process, including review, editing, and final approval of the published version.

## Acknowledgments

We gratefully acknowledge Elettra and XRD2 beamline for providing beamtime and support under proposal 0220565.

## Data availability

The X-ray structure was made available on PDB (PDB ID 7QF6 and 8KD8). The SAXS data and models were deposited to SASbdb under accession codes XXX, YYY, and ZZZ. Primary data were deposited to Zenodo (DOI …, …, …).

## Appendix A. Supplementary data

Supplementary data to this article can be found online at

## Abbreviations

AcCoA: acetyl-CoA
Blast: Basic Local Alignment Search Tool
BME: β-mercaptoethanol
CoA: coenzyme A
CSS: Complex Formation Significance Score
D_max_: maximum dimension
DTNB: 5,5’-dithiobis (2-nitrobenzoic acid)
EC number: Enzyme Commission number
EDTA: Ethylenediaminetetraacetic acid
FC: ferricrocin
FPLC: Fast protein liquid chromatography
FsC: fusarinine C
GNAT: GCN5-related N-acetyltransferase
GSSH: oxidized glutathione
NAD: nicotinamide adenine dinucleotide
NCBI: National Center for Biotechnology Information
NRPS: nonribosomal peptide synthetase
NSD: normal spatial discrepancy
P(r): pair-distribution function
PDB: Protein Data Bank
PEG: polyethylene glycol
PISA: Protein Interfaces, Surfaces and Assemblies
RCSB: Research Collaboratory for Structural Bioinformatics
Rg: radius of gyration
RIA: reductive iron assimilation RMSD root mean square
SAXS: Small Angle X-ray Scattering
SEC: size-exclusion chromatography
SIA: Siderophore-mediated Iron Acquisition
TAFC: triacetylfusarinine C
TM-score: Template Modeling score

