## supplementary material for "Structural and functional characterization of SidF, a possible dual substrate *Aspergillus fumigatus* N5-acetyl-N5-hydroxy-L-ornithine transacetylase"

**Supplement table**

**Table S1** Mutagenesis primer sequences

**Table S2** Essential SAXS data acquisition, sample details, data analysis, modelling fitting and software used

**Table S1** Mutagenesis primer sequences

| Name | Sequence 5' to 3' |
|---|---|
| Fw_NterDeletion | CGGCCGATTTGGGCGGTG |
| Fw_NterDel2 | ACTATGAGTTTAGCATGAAATTCCCGT |
| Fw_NterDel3 | AGCTTGGTCTACAGCCGTT |
| Rv_NterDeletion | ATGGCCCTGAAAATAAAGATTCTCGCC |
| Fw_CterDeletion | TAAGGATCCGAATTCGAGCTCCGTC |
| Rv_CterDeletion | CGGGCCCACCGGACTACCt |
| Fw-Ctrim | TAAGGATCCGAATTCGAGC |
| Rv-Ctrim | CCAGAACAGCGGGCATAA |

**Table S2** Essential SAXS data acquisition, sample details, data analysis, modelling fitting and software used

| **(*a*) Sample details** | | | |
|---|---|---|---|
| | SidF | SidF C-trim | SidF N-terminal domain |
| SASbdb ID | | | |
| Organism | *A. fumigatus* | *A. fumigatus* | *A. fumigatus* |
| UniProt sequence ID (residues in construct) | Q4WF55 (1-462) | Q4WF55 (1-443) | Q4WF55 (1-199) |
| Extinction coefficient [A280, 0.1%(w/v)] | 2.074 | 2.057 | 1.484 |
| Partial specific volume $\bar{\upsilon}$ ($cm^3 g^{-1}$) | 0.73 | 0.731 | 0.731 |
| Particle contrast from sequence and solvent constituents, $\Delta\bar{\rho}$($\rho$ protein - $\rho$ solvent; $10^{10} cm^{-2}$) | 2.911 (12.388 - 9.478) | 2.900 (12.378 - 9.478) | 2.908 (12.386 - 9.478) |
| Molecular mass of monomer *M* from chemical composition (Da) | 55710 | 53496 | 24551 |
| For SEC-SAS, loading volume/concentration, (mg $ml^{-1}$)<br>injection volume (µl), flow rate (ml $min^{-1}$) | 12.7 mg/ml, 400 µl, 0.05 ml $min^{-1}$ | - | - |
| Concentration (range/values) measured and method | | 5 mg/ml | 5 mg/ml |
| Solvent composition and source | 50 mM Tris, 200 mM NaCl pH 8.0 | | |

| **(*b*) SAXS data collection parameters** | | | |
|---|---|---|---|
| Source, instrument and description or reference | SAXSpoint 2.0, Anton Paar with MetalJet C2 and Eiger 1M | | |
| Wavelength (Å) | 1.34 | | |
| Beam geometry (size, sample-to-detector distance) | | | |
| Beam size ($mm^2$) | 1x1 | 1x1 | 1x1 |
| Sample to detector distance (mm) | 792.6 | 571.0 | 571.0 |
| *q*-measurement range ($Å^{-1}$ ) | *0.0068 – 0.43* | *0.008 – 0.59* | *0.008 – 0.59* |
| Basis for normalization to constant counts | Semi-transparent beamstop | | |
| Method for monitoring radiation damage, X-ray dose where relevant | Correlation between individual frames | | |
| Exposure time, number of exposures | | 60x15s | 60x15s |
| Sample configuration including path length and flow rate where relevant | Flow through quartz kapillary (2mm) with UV-Vis absroption posibility | Quartz kapillary (1.5mm) | |
| Sample temperature (ºC) | 20 | | |

| **(*c*) Software employed for SAXS data reduction, analysis and interpretation** | |
|---|---|
| SAXS data reduction to sample–solvent scattering, and extrapolation, merging, desmearing *etc*. as relevant | Reduction: in-house solution<br>solvent subtraction: PRIMUSqt [1]<br>SEC-SAXS chromatograph: RAW [2] |
| Calculation of ε from sequence | ProtParam [3] |
| Calculation of $\Delta\bar{\rho}$ and $\bar{\upsilon}$ values from chemical composition | MULCh 1.1 [4] |
| Basic analyses: Guinier, *P*(*r*), scattering particle volume (*e.g.* Porod volume $V_P$ or volume of correlation $V_c$) | PRIMUSqt from ATSAS 3.2.1<br>RAW |
| Shape/bead modelling | DAMMIF, DAMMIN , DAMAVER |
| Atomic structure modelling (homology, rigid body, ensemble) | CRYSOL from PRIMUSqt in ATSAS 3.2.1, EOM [5], CORAL [6] |
| Molecular graphics | PyMOL v.2.5.5 Win64<br>Chimera |

| **(*d*) Structural parameters** | | | |
|---|---|---|---|
| Guinier Analysis | SidF | SidF C-trim | SidF N-terminal domain |
| *I*(0) (A.U.) | 0.85 +/- $3.81 \times 10^{-3}$ | 0.28 +/- $1.09 \times 10^{-3}$ | 0.04 +/- $3.79 \times 10^{-4}$ |
| $R_g$ (Å) | 40.49 +/- 0.26 | 34.52 +/- 0.2 | 21.85 +/- 0.33 |
| *q*-range (Å$^{-1}$) | 0.0112 - 0.0325 | 0.0129 - 0.0373 | 0.0147 - 0.0582 |
| Coefficient of correlation, $R^2$ | 0.994 | 0.995 | 0.925 |
| *M* from Bayesian Inference (Da) | 208,000 | 109,100 | 21,200 |
| *P*(*r*) analysis | SidF | SidF C-trim | SidF N-terminal domain |
| *I*(0) (A.U.) | 0.87 +/- $4.87 \times 10^{-3}$ | 0.28 +/- $1.66 \times 10^{-3}$ | 0.04 +/- $4.29 \times 10^{-4}$ |
| $R_g$ (Å) | 43.2 +/- 0.48 | 36.81 +/- 0.39 | 23.24 +/- 0.4 |
| $d_{max}$ (Å) | 190 | 150 | 90 |
| *q*-range (Å$^{-1}$) | 0.0112 - 0.3997 | 0.0129 - 0.3997 | 0.0147 – 0.3997 |
| $X^2$ (total estimate from GNOM) | 1.285 | 1.017 | 1.133 |
| Volume (*e.g.* $V_P$ and/or $V_c$) | $2.94 \times 10^5$, 1.01E+03 | $1.44 \times 10^5$, | $2.85 \times 10^4$, 241.4 |

| ***e1*) Atomistic modelling** | | | | | | |
|---|---|---|---|---|---|---|
| Experimental SAXS | **SidF** | **SidF** | **SidF** | **SidF** | **SidF N-ter** | **SidF N-ter** |
| Calculated SAXS | **SidF Monomer** | **SidF Dimer(A:F)** | **SidF Dimer(A:D)** | **SidF Tetramer** | **SidF N-ter Monomer** | **SidF N-ter Dimer** |
| **q range for all modelling** | 0.0089 - 0.3997 | | | | 0.01 – 0.3997 | |
| ***CRYSOL*** (with default parameters and constant subtraction allowed) | | | | | | |
| $X^2$, P-value | 79.7, 0.0 | 9.624, 0.0 | 9.114, 0.0 | 2.172, 0.00 | 2.116, 0.0 | 5.362, 0.0 |
| Predicted Rg (Å) Rg from the slope of net intensity [A] | 24.86 | 30.01 | 24.86 | 35.92 | 20.53 | 26.41 |
| Ex. Vol (Å$^3$) | 63,205 | 125,390 | 126,310 | 253,040 | 24,510 | 48,557 |
| ***FoXS*** (with default parameters) | | | | | | |
| $X^2$, P-value | 292.78, 0.00 | 69.66, 0.00 | 75.52, 0.00 | 2.49, 0.00 | 1.98, 0.00 | 5.00, 0.00 |
| Predicted Rg (Å) | 23.58 | 28.71 | 30.80 | 34.95 | 20.70 | 25.59 |
| $c_1$, $c_2$ | 1.05, 4.00 | 1.05, 4.00 | 1.05, 4.00 | 1.02, 4.00 | 0.99, 4.00 | 1.03, -2.00 |
| ***EOM*** | | | | | | |
| $X^2$ | - | - | - | 1.1 | 1.188 | |
| Predicted Rg (Å) | - | - | - | 40.71 | 22.7 | |

| *e2*) **Atomistic modelling** | | | | |
|---|---|---|---|---|
| Experimental SAXS | **SidF C-trim** | **SidF C-trim** | **SidF C-trim** | **SidF C-trim** |
| Calculated SAXS | **SidF Monomer** | **SidF Dimer(A:F)** | **SidF Dimer(A:D)** | **SidF Tetramer** |
| **q range for all modelling** | 0.01 - 0.3997 | | | |
| ***CRYSOL*** (with default parameters and constant subtraction allowed) | | | | |
| $X^2$, P-value | 12.03, 0.0 | 9.906, 0.0 | 6.918, 0.0 | 142.9, 0.00 |
| Predicted Rg (Å) Rg from the slope of net intensity [A] | 24.86 | 30.01 | 31.68 | 35.92 |
| Ex. Vol (Å$^3$ ) | 63,205 | 125,390 | 126,310 | 253,040 |
| ***FoXS*** (with default parameters) | | | | |
| $X^2$, P-value | 64.72, 0.00 | 10.59, 0.00 | 10.22, 0.00 | 95.33, 0.00 |
| Predicted Rg (Å) | 23.58 | 28.71 | 30.80 | 34.95 |
| $c_1$, $c_2$ | 1.05, 4.00 | 1.01, -0.41 | 1.02, 0.91 | 1.05, -2.00 |
| ***CORAL*** *(N and C terminal domain were modelled separately, no chain specific)* | | | | |
| $X^2$ | | 1.87 | | |
| Predicted Rg (Å) | | 34.92 | | |

**Supplementary Figures**

**Figure S1** Schematic of a typical acyltransferase reaction of SidF.

**Figure S2** SAXS data summary for SidF full-length. (A) Scattering profile on a log-lin scale. (B) Guinier fit (top) and fit residuals (bottom). (C) Dimensionless Kratky plot. Dashed lines show where a globular system would peak. (D) P(r) functions, normalized by I(0). Orange is P(r) truncated for DAMIF.

**Figure S3** The SidF tetramer X-ray structure dimension calculated in Pymol (top) give the size of 98.7 x 86.8 x 79.8 Å.

**Figure S4** SidF protein variant constructions. The black solid line represents the missing part compared to wild-type protein.

**Figure S5** SAXS data summary for SidF N-terminal domain. (A) Scattering profile on a log-lin scale. (B) Guinier fit (top) and fit residuals (bottom). (C) Dimensionless Kratky plot. Dashed lines show where a globular system would peak. (D) P(r) functions, normalized by I(0). Orange is P(r) truncated for DAMIFF.

**Figure S6** SAXS data summary for SidF C-trim. (A) Scattering profile on a log-lin scale. (B) Guinier fit (top) and fit residuals (bottom). (C) Dimensionless Kratky plot. Dashed lines show where a globular system would peak. (D) P(r) functions, normalized by I(0).

**Figure S7** Melting temperature (Tm) of SidF variants analyzed by DSF

(Left) Normalized averaged raw data for SidF full-length (WT), N-terminal domain (N), and C-terminus (C-trim) are shown. The averaged Tm values are presented with standard deviation (n=6) and demonstrated in bar chart (right). P-values indicate the statistical significance of the ΔTm between SidF full-length and the N-terminal domain (N) or C-terminus (C-trim). A P-value < 0.0005 is denoted by three asterisks (***).

**Figure S8** The effect of AcCoA and hydroxy-L-ornithine (OH-L-orn) on SidF full-length melting temperature (Tm) determined by DSF.

Normalized averaged raw data for SidF full-length incubated with AcCoA, OH-L-orn, and a combination of both (AcCoA + OH-L-orn) are shown. The averaged Tm values for each condition are presented with standard deviation (n = 6).

**Figure S9** Melting temperature (Tm) of SidF variants analyzed by nanoDSF

(A) Normalized raw data for SidF full-length (WT), (B) N-terminal domain, and (C) C-terminus are shown. (D) The averaged Tm values are presented with standard deviation and demonstrated in bar chart. P-values indicate the statistical significance of the ΔTm between SidF full-length and the N-terminal domain or C-terminus. A P-value < 0.0005 is denoted by three asterisks (***). (∗∗∗$p < 0.0005$; ∗∗$p < 0.005$; ∗$p < 0.05$)

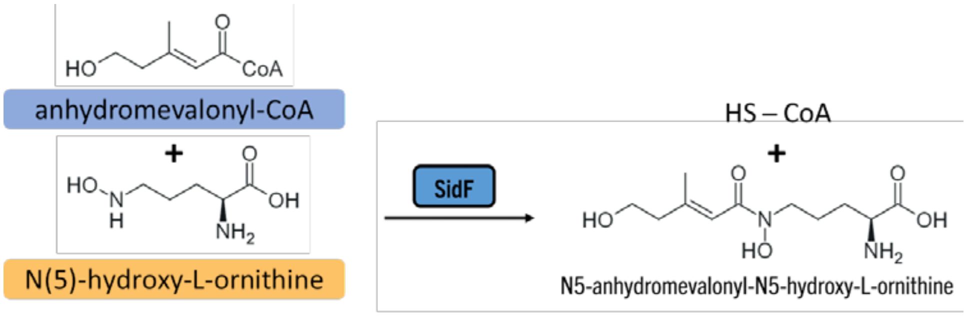


Figure S1

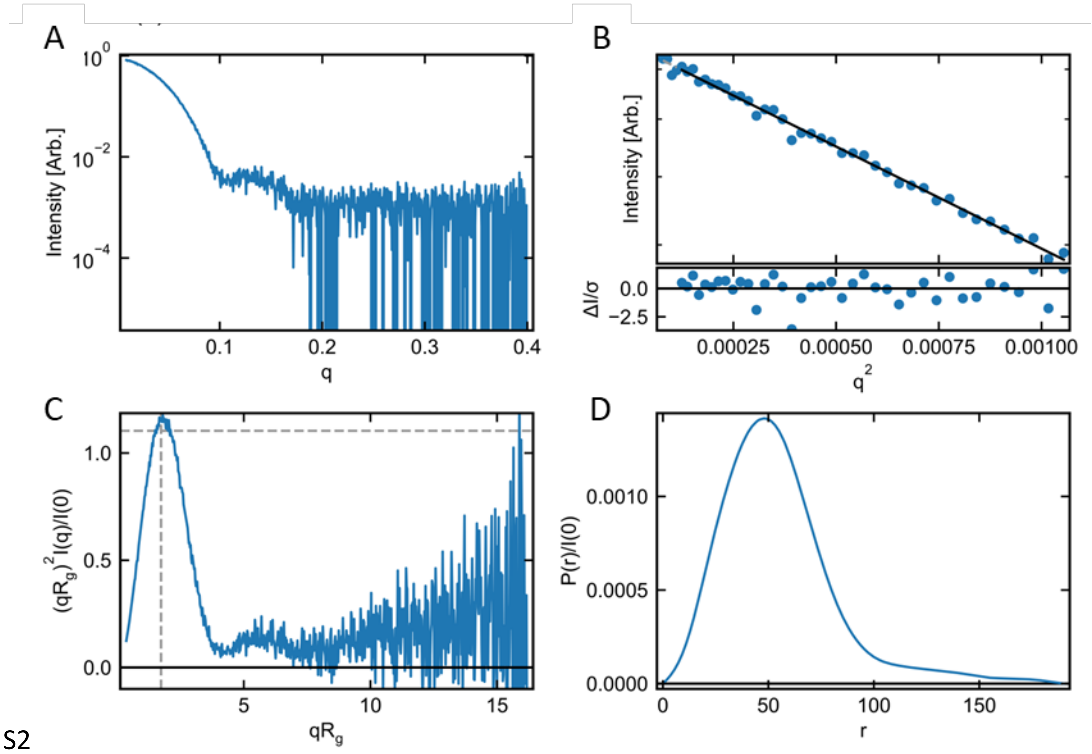


Figure S2

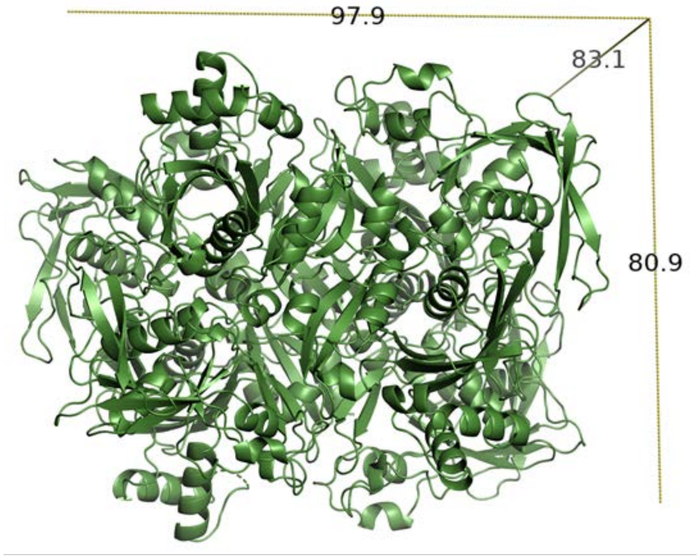


Figure S3

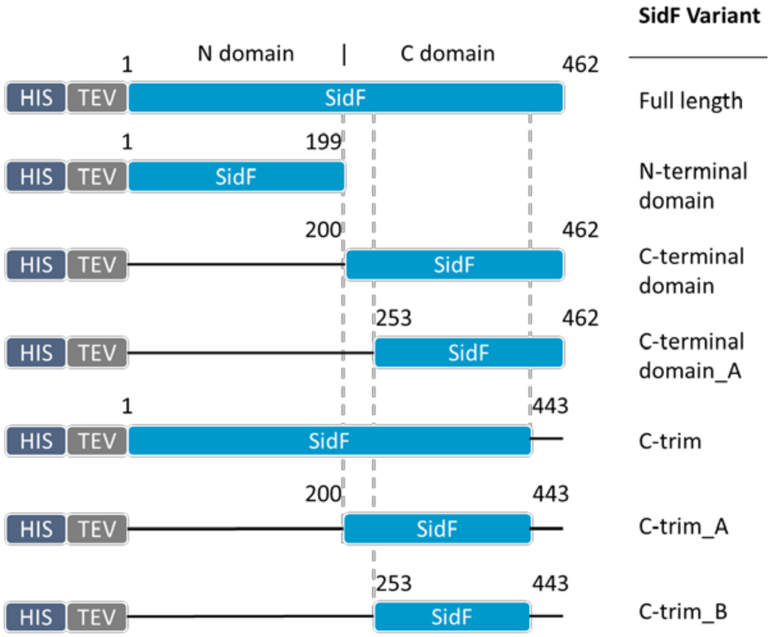


Figure S4

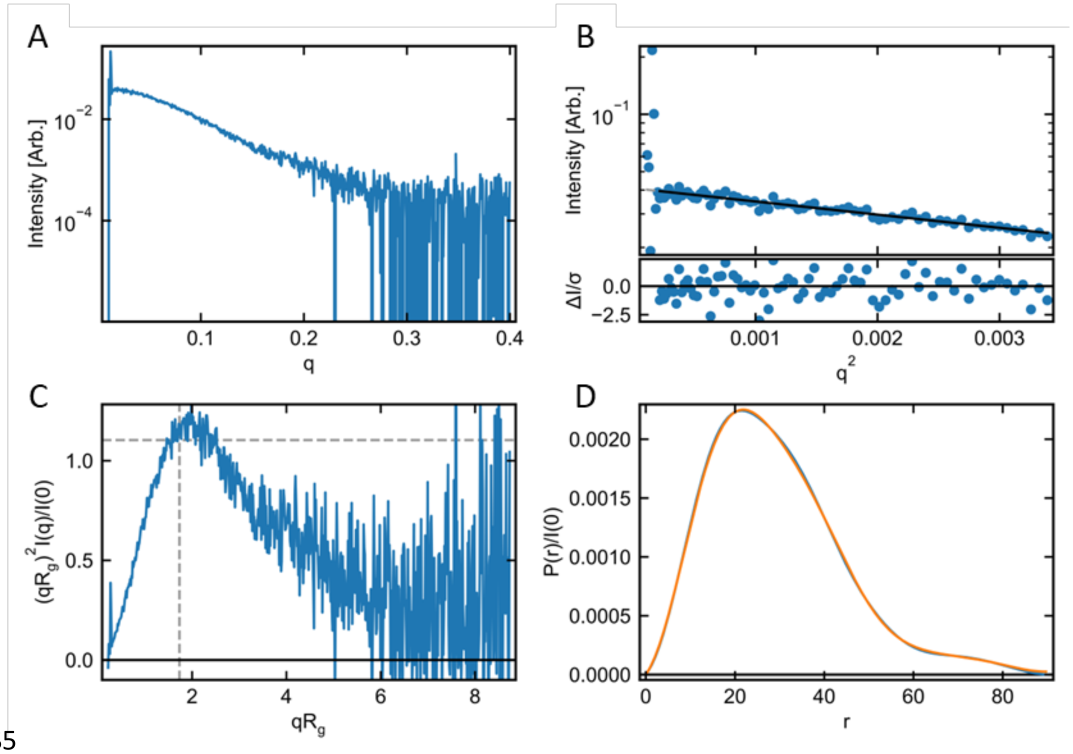

Figure S5

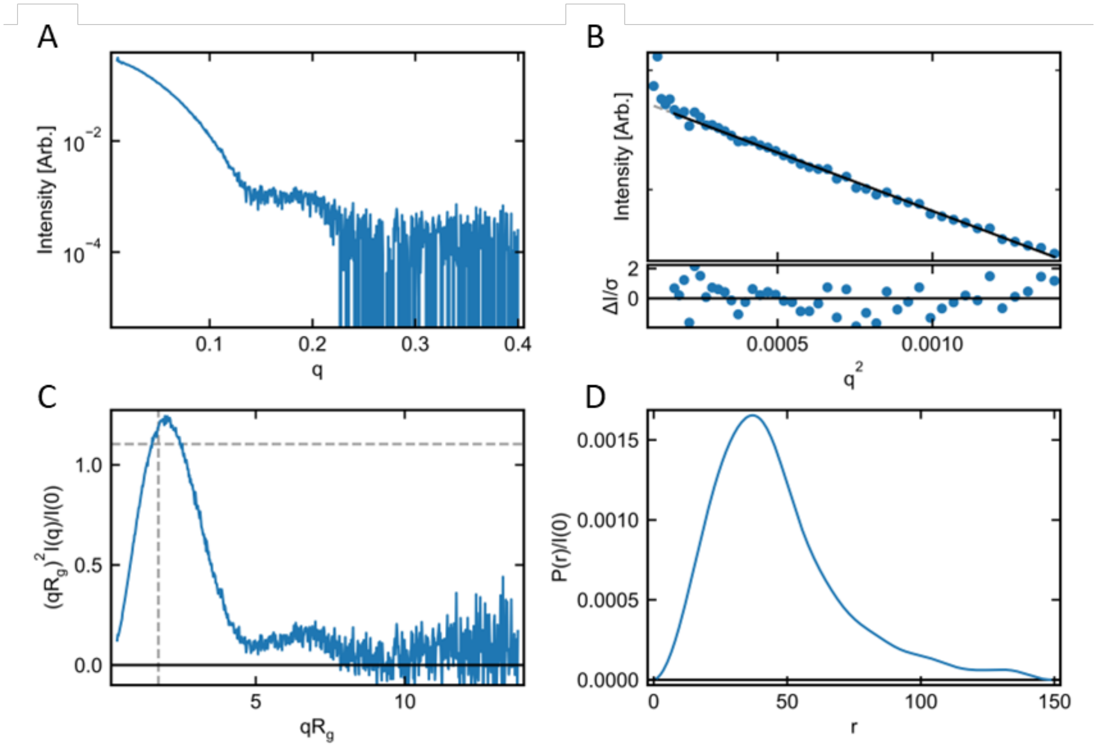

Figure S6

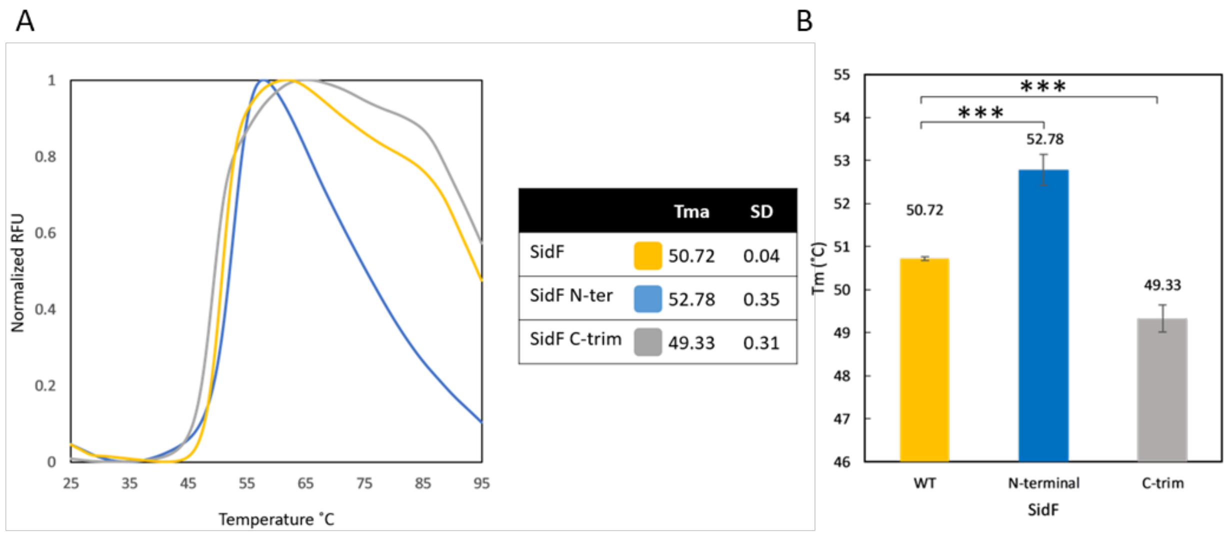
A
Normalized RFU
1
0.8
0.6
0.4
0.2
0
25
35
45
55
65
75
85
95
Temperature ˚C
Tma
SD
SidF
50.72
0.04
SidF N-ter
52.78
0.35
SidF C-trim
49.33
0.31
B
Tm (˚C)
55
54
53
52
51
50
49
48
47
46
***
***
50.72
52.78
49.33
WT
N-terminal
C-trim
SidF

Figure S7

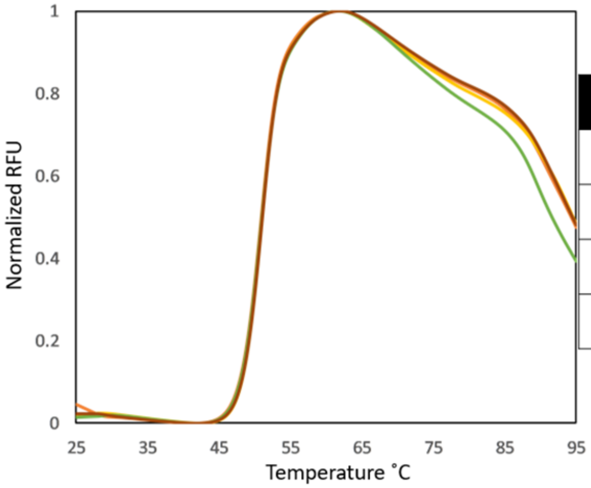


| | Tma | SD |
|---|---|---|
| SidF | 50.72 | 0.04 |
| SidF+AcCoA | 50.8 | 0.11 |
| SidF+OH-L-Orn | 50.73 | 0.1 |
| SidF+AcCoA+OH-L-Orn | 50.85 | 0.08 |

Figure S8

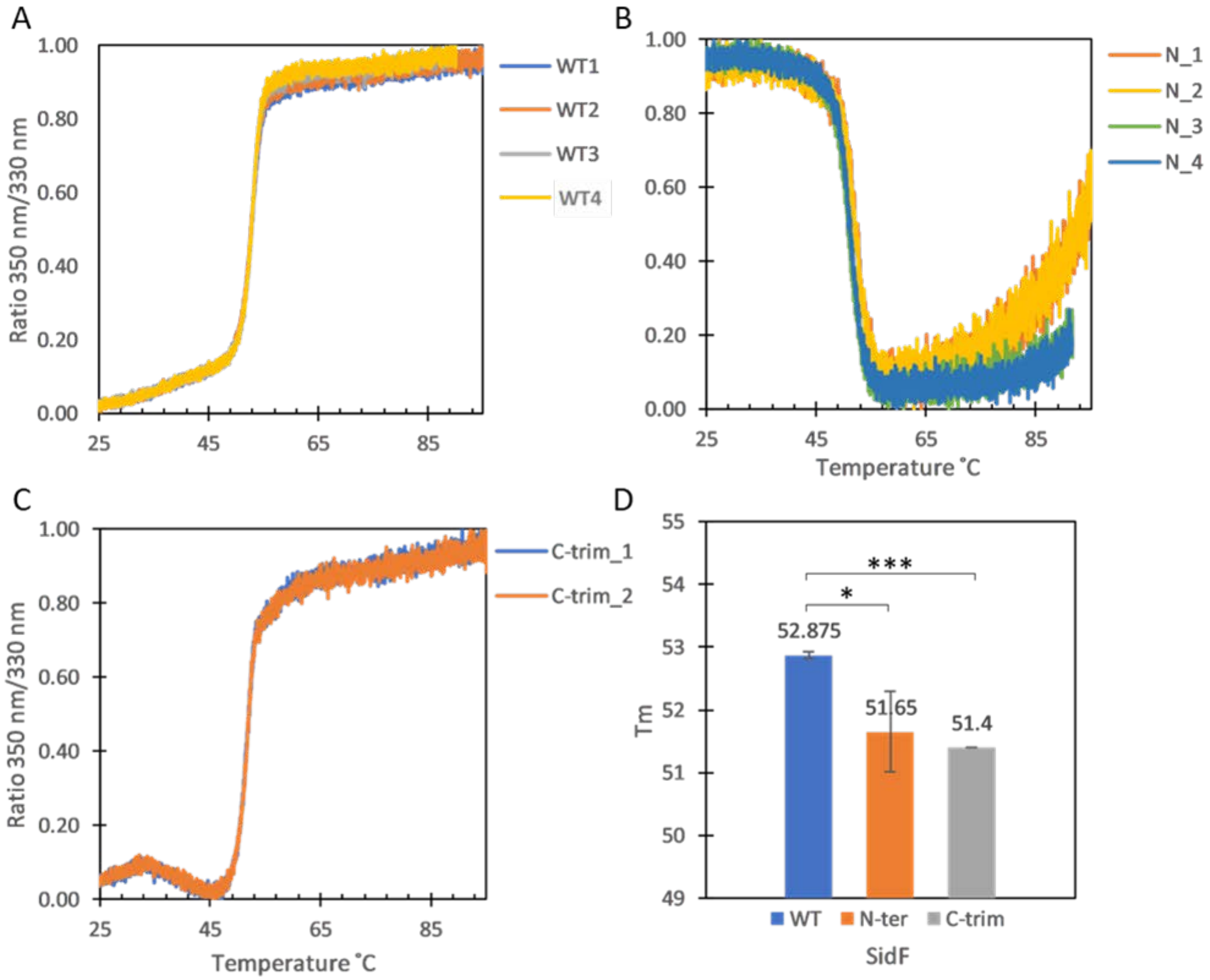

A
1.00
0.80
0.60
0.40
0.20
0.00
Ratio 350 nm/330 nm
25
45
65
85
WT1
WT2
WT3
WT4
B
N_1
N_2
N_3
N_4
Temperature ˚C
C
C-trim_1
C-trim_2
D
55
54
53
52
51
50
49
Tm
***
*
52.875
51.65
51.4
WT
N-ter
C-trim
SidF


Figure S9

**Thermal Shift Assay For Protein Stability Examination**

The 20 µl reactions containing protein and ProteOrange 5000X (Lumiprobe) at final concentrations of 5 µM and 5x, respectively, supplemented with or without ligands at final concentrations of 25 µM were set up in 96-well PCR plates. The reactions were equilibrated at 25°C for 2 min, then increased to 95°C at a rate of 2°C min$^{-1}$. The experiments were performed using a FRET channel in Bio-Rad CX1000 Real-Time PCR Systems (Bio-Rad CFX Maestro 1.1 software version 4.1.2433.1219) in two independent replicates (total n = 6 for each sample). The melting temperature (Tm) was determined from the first derivative using DSFworld [7].

**Analysis of SidF variant protein stability using DSF assay**

The stability of SidF full-length, N-terminal domain, and C-trim was investigated by protein thermal shift assay. The stability of the N-terminal domain increased when compared to the WT protein with the melting temperature difference (ΔTm) of 1.95℃ (p-value = 0.000059) (Fig. S11). In contrast, the SidF C-trim stability was decreased compared to the WT protein with the ΔTm of -1.5℃ (p-value = 0.000372). SAXS analysis revealed that SidF full-length exists as a tetramer, while the N-terminal domain adopts a monomeric state and the C-trim forms a dimer. Potential explanations for the observed stability variations can be attributed to the differences in oligomeric state. The N-terminal domain, existing as a monomer, possesses a single domain structure, inherently less flexible compared to the full-length protein which has two distinct domains. Also, it is potentially more stable than the complex comprising four subunits, where there may still be some degree of movement between each domain and each subunit. This could potentially lead to a more rigid structure and increased thermal stability. The tetrameric form, presumed to represent its biological configuration, is expected to adopt its lowest energy state. Stabilization of the tetramer occurs through hydrogen bonds within the dimer as well as between dimers. The C-trim exists as a dimer, lacking the inter-dimer interactions present in the full-length tetramer. This potentially allows for increased flexibility and movement within the C-trim compared to the full-length protein, resulting in decreased thermal stability.

The thermal shift assay did not detect a significant change (Tm shift) upon incubation of SidF protein with its substrates (Fig. S12). However, enzymatic activity assays confirmed the occurrence of the reaction, implying ligand binding. This suggests that substrate binding may not induce a substantial alteration in SidF protein stability, potentially rendering the thermal shift assay insensitive for detecting this interaction.

**NanoDSF for Protein Stability Examination**

The SidF WT and C-trim proteins were diluted to the concentrations of 0.5 mg/ml (8.98 µM), while SidF N-terminal domain was diluted to 1 mg/ml (40 µM). The samples were loaded into Prometheus NT.48 nanoDSF Grade Standard Capillaries (NanoTemper Technologies). The reactions were equilibrated at 20°C followed by an increase to 95°C at 1.5°C /min. The experiments were performed using Prometheus NT.48 (NanoTemper Technologies) at least in two replicates. The Tm was determined from the first derivative of the 350 nm/330 nm ratio.

**Analysis of SidF Variant Protein Stability using nanoDSF Assay**

The thermal stability of full-length SidF protein, its N-terminal domain (N-ter), and C-terminal truncation (C-trim) was investigated using nanoDSF. Compared to the wild-type (WT) protein, the N-ter and C-trim exhibited decreased stability, reflected by a melting temperature difference (ΔTm) of 1.23 °C (p-value = 0.0088156) and 1.48 °C (p-value = 0.0000025), respectively (Fig. S13).

Interestingly, during unfolding, the N-ter displayed a blue shift in its fluorescence spectrum (inverted peak with stronger decrease at 350 nm compared to 330 nm), resulting in a lower fluorescence intensity ratio (350 nm/330 nm). This phenomenon may be attributed to the surface-exposed tryptophan (Trp) residues in the N-ter. These residues might be readily accessible at the beginning of the experiment, leading to a high initial fluorescence at 350 nm. Subsequently, as temperature increases, Trp residues become buried within the denatured protein's hydrophobic environment, leading to a decreased fluorescence signal.

We observed significant differences (p-value < 0.0005 for WT, p-value = 0.0066 for N-ter, p-value = 0.0001 for C-trim) in Tm values obtained from nanoDSF compared to conventional DSF. Notably, the C-trim Tm from both methods followed the same trend, showing a decrease compared to the WT. However, the N-ter displayed contrasting results – increased Tm with conventional DSF and decreased Tm with nanoDSF.

These discrepancies likely arise from the fundamental differences in the measurement principles of each technique. Conventional DSF relies on the binding of a hydrophobic fluorescent dye to the exposed hydrophobic regions in the unfolding protein. In contrast, nanoDSF utilizes the intrinsic fluorescence signal of Trp and tyrosine (Tyr) residues, measured at 350 nm and 330 nm. Dye-protein interactions in conventional DSF can potentially influence the measured Tm, introducing artifacts. Additionally, it might exhibit reduced sensitivity for smaller proteins like truncated variants due to weaker dye binding. Similarly, nanoDSF has limitations when comparing proteins of different sizes. The presence and location of Trp and Tyr residues can vary in truncated proteins, affecting the fluorescence signal strength and interpretation.

Both DSF and nanoDSF present limitations when directly comparing the stability of proteins with significant size variations. We acknowledge these limitations and focus on qualitative trends rather than absolute Tm values. To obtain a more comprehensive assessment of protein stability, complementary techniques like Circular Dichroism (CD) or Differential Scanning Calorimetry (DSC) should be introduced.
